# gapseq: Informed prediction of bacterial metabolic pathways and reconstruction of accurate metabolic models

**DOI:** 10.1101/2020.03.20.000737

**Authors:** Johannes Zimmermann, Christoph Kaleta, Silvio Waschina

## Abstract

Microbial metabolic processes greatly impact ecosystem functioning and the physiology of multi-cellular host organisms. The inference of metabolic capabilities and phenotypes from genome sequences with the help of reference biomolecular knowledge stored in online databases remains a major challenge in systems biology. Here, we present `gapseq`: a novel tool for automated pathway prediction and metabolic network reconstruction from microbial genome sequences. `gapseq` combines databases of reference protein sequences (UniProt, TCDB), in tandem with pathway and reaction databases (MetaCyc, KEGG, ModelSEED). This enables the prediction of an organism’s metabolic capabilities from sequence homology and pathway topology criteria. By incorporating a novel LP-based gap-filling algorithm, `gapseq` facilitates the construction of genome-scale metabolic models that are suitable for metabolic phenotype predictions by using constraint-based flux analysis. We validated `gapseq` by comparing predictions to experimental data for more than 3, 000 bacterial organisms comprising 14, 895 phenotypic traits that include enzyme activity, energy sources, fermentation products, and gene essentiality. This large-scale phenotypic trait prediction test showed, that `gapseq` yields an overall accuracy of 81% and thereby outperforms other commonly used reconstruction tools. Furthermore, we illustrate the application of `gapseq`-reconstructed models to simulate biochemical interactions between microorganisms in multi-species communities. Altogether, `gapseq` is a new method that improves the predictive potential of automated metabolic network reconstructions and further increases their applicability in biotechnological, ecological, and medical research. `gapseq` is available at https://github.com/jotech/gapseq.

## 1 Background

Anything you have to do repeatedly may be ripe for automation.

— Doug McIlroy

Metabolism is central for organismal life. It provides metabolites and energy for all cellular processes. A majority of metabolic reactions are catalysed by enzymes, which are encoded in the genome of the respective organism. Those catalysed reactions form a complex metabolic network of numerous biochemical transformations, which the organism is presumably able to perform [1].

In systems biology, the reconstruction of metabolic networks plays an essential role, as the network represents an organism’s capabilities to interact with its biotic and abiotic environment and to transform nutrients into biomass. Mathematical analysis has shown great potential for dissecting the functioning of metabolic networks on the level of topological, stoichiometric, and kinetic models [2], which together provide a wide array of methods [3]. Although different microbial metabolic modelling approaches exist, they can be summarised by a theoretical framework that provides a unifying view on microbial growth [4]. Metabolic models not only have demonstrated their ability to predict phenotypes on the level of cellular growth and gene knockouts, but also provide potential molecular mechanisms in form of gene and reaction activities, which can be validated experimentally [5]. Due to this predictive potential, genome-scale metabolic models have been applied to identify metabolic interactions between different organisms [6, 7, 8, 9, 10], to study host-microbiome interactions [11, 12, 13], to predict novel drug targets to fight microbial pathogens [14, 15], and for the rational design of microbial genotypes and growth-media conditions for the industrial production or degradation of biochemicals [16, 17]. Furthermore, recent advances in DNA-sequencing technologies have led to a vast increase in available genomic- and metagenomic sequences in databases [18], which further expands the applicability of genome-scale metabolic network reconstructions.

The reconstruction of metabolic networks links genomic content with biochemical reactions and therefore depends on sequence annotations and reaction databases, which are both crucial for overall network quality [19, 20]. A general problem in reconstructing metabolic networks occurs by an incorrect representation of the organism’s physiology. First, inconsistencies in databases can lead to an incorporation of imbalanced reactions into the metabolic network, which may become responsible for incorrect energy production by futile cycles [20]. Second, many genes are lacking a functional annotation due to a lack of knowledge [21] and, thus, also the gene products cannot be integrated into the metabolic networks, which potentially lead to gaps in pathways. Third, the gap-filling of metabolic networks is frequently done by adding a minimum number of reactions from a reference database that facilitate growth under a chemically defined growth medium [22, 23, 24]. Such approaches miss further evidences potentially hidden in sequences and are biased towards the growth medium used for gap-filling. And fourth, the validation of predictions made by metabolic networks is so far only performed with smaller experimental data sets from model laboratory strains such as *Escherichia coli* K12 or *Bacillus subtilis* 168 and therefore the overall performance of many metabolic models is insufficiently assured.

Genome-scale metabolic network reconstructions are increasingly applied to simulate complex metabolic processes in microbial communities [25]. Such simulations are highly sensitive to the quality of the individual metabolic networks of the community members. This is because the accurate prediction of fermentation products and carbon source utilisation is crucial for the correct prediction of metabolic interactions since the substances produced by one organism may serve as resource for others [26]. Thus, in multi-species communities, the metabolic fluxes of organisms are intrinsically connected, which can lead to error propagation when one defective model affects otherwise correctly working models. As a consequence, the feasibility of community modeling intrinsically depends on the accuracy of the individual organismal models.

In this work, we present `gapseq` a novel software for pathway analysis and metabolic network reconstruction. The pathway prediction is based on multiple biochemistry databases that comprise information on pathway structures, the pathways’ key enzymes, and reaction stoichiometries. Moreover, `gapseq` constructs genome-scale metabolic models that enable metabolic phenotype predictions as well as the application in simulations of community metabolism. Models are constructed using a manually curated reaction database that is free of energy-generating thermodynamically infeasible reaction cycles. As input, `gapseq` takes the organism’s genome sequence in FASTA format, without the need for an additional annotation file. Topology as well as sequence homology to reference proteins inform the filling of network gaps, and the screening for potential carbon sources and metabolic products is done in a way that reduces the impact of growth medium definitions. Finally, we used large-scale experimental data sets to validate enzyme activity, carbon source utilisation, fermentation products, gene essentiality, and metabolite-cross feeding interactions in microbial communities.

## 2 Results

### 2.1 Biochemistry database and universal model

The pathway-, transporter, and complex prediction is based on a protein sequence database that is derived from UniProt as well as TCDB and consists in total of 130,671 unique sequences (111,542 reviewed unipac 0.9 clusters and 19,129 TCDB transporter) and also 1,131,132 unreviewed unipac 0.5 cluster that can be included optionally. In addition, the protein sequence database in `gapseq` can be updated to include new sequences from Uniprot and TCDB. For the construction of genome-scale metabolic network models we have built a biochemistry database, that is derived from the ModelSEED biochemistry database. In total, the resulting curated `gapseq` metabolism database comprises 14.287 reactions (including transporters) and 7.570 metabolites. All metabolites and reactions from the biochemistry database are incorporated in the universal model that `gapseq` utilises for the gap-filling algorithm. When removing all dead-end metabolites and corresponding reactions, the universal model comprises 10.194 reactions and 3.337 metabolites. It needs to be noted, that the current biochemistry database and the derived universal model represents bacterial metabolic functions and that, at the current version of `gapseq`, the database does not include archaea-specific reactions. However, those reactions and, thus, also the possibility to use `gapseq` for the reconstruction of archaeal models will be included in an later version of the software.

### 2.2 Agreement with enzymatic data (BacDive)

We used experimental data of active metabolic enzymes to compare the accuracy of model generation pipelines. In total, we compared 10,538 enzyme activities, comprising 30 unique enzymes, in 3,017 organisms. For all organisms, genome-scale metabolic models were constructed using three different pipelines (CarveMe[39], `gapseq`, ModelSEED[24]). `gapseq` models had with 6% the lowest false-negative rate compared to CarveMe (32%) and ModelSEED (28%). Correspondingly, `gapseq` showed with 53% also highest true positive rate compared to CarveMe (27%) and ModelSEED (30%), while the rates of false positive and true negative predictions were comparable (Figure 1A). For this test, the most prominent EC numbers were the catalase, 1.11.1.6, accounting for 26% of the comparisons and the cytochrome oxidase, 1.9.3.1, accounting for 22%.

**Figure 1:**
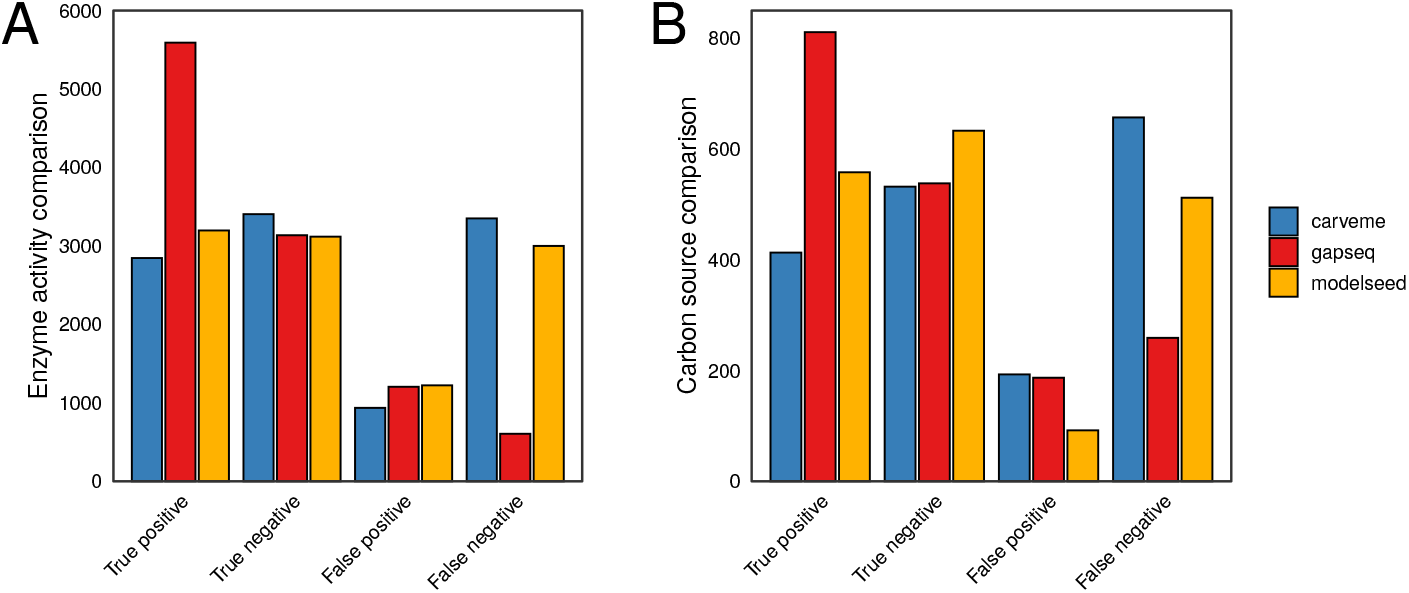
Results from enzyme activity and carbon source validations. A) In total 10,538 enzyme activities (30 enzymes and 3,017 organisms) from experimental standardised experiments from the DSMZ BacDive database were compared for three different methods. B) The predictions of 1,795 carbon sources (48 unique carbon sources and 526 organisms) were validated with data from the ProTraits database.

### 2.3 Validation of carbon source usage (ProTraits)

Growth predictions are essential for metabolic models. We checked the quality of model generation pipelines to predict the growth on different carbon sources. In summary, we compared 1,795 different growth prediction for 526 organism and 48 carbon sources (Figure 1B). `gapseq` outperformed the other methods in terms of false negatives (14% compared with 29% ModelSEED and 37% CarveMe) and true positives (45% compared with 31% ModelSEED and 23% CarveMe). ModelSEED showed fewer false positives (5% compared with 10% `gapseq` and 11% CarveMe) and more true negatives (35% compared with 30% `gapseq` and 30% CarveMe). `gapseq`, predicted most false positives for formate (29 times). This overestimate of formate as potential carbon source is likely due to the fact that we tested carbon source utilisation on the basis of electron transfer from the source to electron carriers (i.e. ubiquinol, menaquinol, or NADH), which is analogous to the experiemental carbon source test of BIOLOG plates [46]. However, while it is known that formate can serve in fact as electron donor in a number of different bacteria [84], the role as source of carbon atoms for the synthesis of biomass components is limited to a few known methylotrophs [85].

Across all methods, the most accurately predicted carbon sources, with more than 100 tested organisms, were fructose (91% correct predictions), mannose (89%), or arginine (84%), whereby less good predictions were obtained for arabinose (29% correct predictions), dextrin (40%), or acetate (42%).

### 2.4 Gene essentiality

We compared the ability of `gapseq` models to predict the essentially of genes with predictions from ModelSEED and CarveMe reconstructions as well as with curated models for the same organisms (Figure 2). As expected, the curated models outperform all three automated reconstruction tools for most species and prediction metrics (namely precision, sensitivity, specificity, accuracy, and F1-score). Interestingly, for *Pseudomonas aeruginosa* the `gapseq` model shows better gene essentiality predictions in terms of sensitivity, accuracy, and F1-score than the curated model (Figure 2D). Compared to CarveMe, `gapseq` shows generally a higher sensitivity in essentiality predictions but, at the same time, a lower precision rate. This pattern is attributed to the fact, that `gapseq` models tend to predict more genes as essential than CarveMe, leading to a higher number of true positive (TP) predictions but also more false positives (FP). For most organisms and on the basis of most prediction metrics, `gapseq` outperforms network models that were reconstructed using ModelSEED.

**Figure 2:**
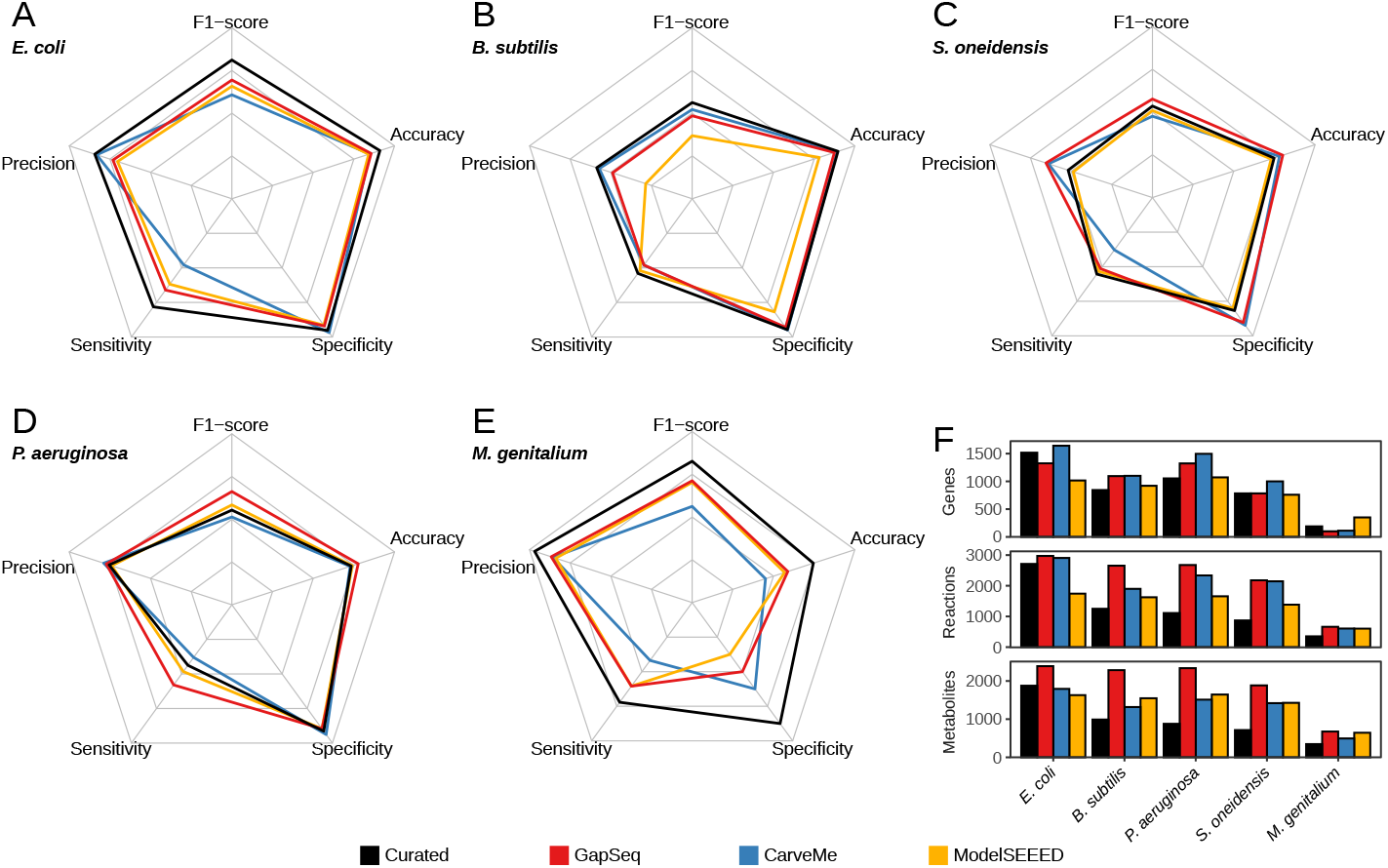
Results from model gene essentiality tests for five bacterial species. (A) *Escherichia coli*, (B) *Bacillus subtilis*, (C) *Shewanella oneidensis*, (D) *Pseudomonas aeruginosa*, and (E) *Mycoplasma genitalium*. Results from `gapseq` models (red) are compared to CarveMe (blue) and ModelSEED (yellow) models, as well as to published curated genome-scale metabolic models (black) of the respective organisms. (F) Counts of genes, reactions (including exchanges and transporters), and metabolites in each reconstruction.

### 2.5 Fermentation products

Anaerobic or facultative anaerobic bacteria utilise different fermentation pathways in order to extract energy from environmental compounds by chemical transformations in the absence of oxygen. We tested if the identity of fermentation products can be predicted by metabolic network model constructions obtained from `gapseq`, CarveMe, and ModelSEED for 18 different bacterial organisms (Figure 2).

The organisms were selected based on following criteria: (1) the organisms have a published RefSeq genome sequence [52], (2) are known anaerobic or facultative anaerobic organisms, and (3) the identity of fermentation products has been experimentally described and reported in primary literature (Suppl. table S2). Overall, `gapseq` showed the highest number of true positive predictions (TP) with 36 TP predicted with the Minimize-Total-Flux (MTF) and 37 TP predicted with Flux-Variability-Analysis (FVA) which is substantially higher compared to CarveMe (8 TP with MTF, 10 TP with FVA) and ModelSEED (1 TP, 3 TP). The production of the short-chain-fatty-acids acetate, butyrate, and propionate was correctly predicted by `gapseq` in 78% of cases and thereby outcompetes CarveMe (9%) and ModelSEED (0%), which did not predict butyrate or propionate production for any organism tested. Moreover, `gapseq` correctly predicted homolactic fermentation by *Lactobacillus delbrueckii* and *Lactobacillus acidophilus*, which is dominated by lactate as fermentation end-product and also predicted heterolactic fermentation by *Bifidobacterium longum*. However, `gapseq` failed to predict lactate production of organisms that utilise different fermentation strategies, which also yield lactate (e.g. mixed-acid fermentation by *Escherichia coli*). Interestingly, the predicted quantities of fermentation product release is higher for true positive than for false negative predictions (Figure 3). This further suggests, that `gapseq` is able to predict the main fermentation products of bacterial organisms during anaerobic growth based on the organism’s genome sequence.

**Figure 3:**
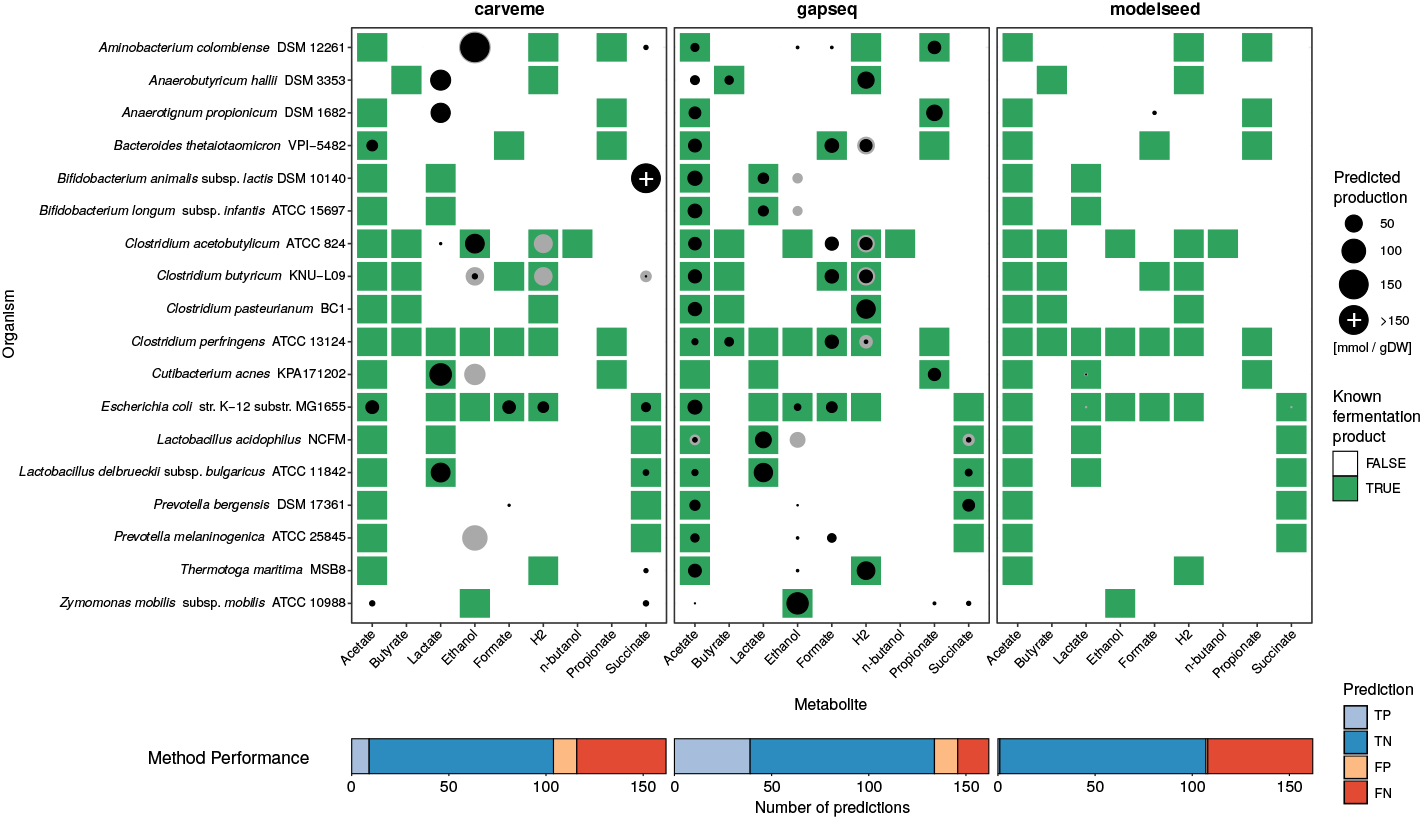
Results of the fermentation product test of 18 bacterial organisms under anaerobic growth with models generated using gapseq, CarveMe, and ModelSEED. Point sizes indicate the predicted production of a fermentation product metabolite (columns) by the corresponding organism (row). Predictions (black) are based on Minimize-Total-Flux (MTF) flux balance analyses. Grey circles indicate the upper production limit obtained from Flux-Variability-Analysis (FVA). Metabolite-organism-combinations highlighted in green denote known fermentation products, which have been reported in literature based on experimental measures of the metabolite in anaerobic cultures.

### 2.6 Anaerobic food web of the gut microbiome

The prediction of metabolic interactions between microbial organisms is of special interest in ecology, medicine, and biotechnology. So far, we showed the capacity of `gapseq` on the level of individual models. In a next step, we simulated several individual models together as a multi-species community to validate the potential of `gapseq` in microbial community modelling. As sample application we selected representative members of the human gut microbiome that are known to form an anaerobic food web [64, 65]. Altogether, we employed 20 organisms and simulated the combined growth in a shared environment for several time steps using the community modeling framework BacArena [68]. On the community level, simulations using `gapseq` models captured all important substances, which are known to be produced in the context of the food web (Figure 4). This included the production of short chain fatty acids (acetate, propionate, butyrate), lactate, hydrogen, hydrogen sulfide (H_2_S), methane, formate, and succinate. The formation of acetate, formate, and hydrogen was most prevalent, which are also common end-products of fermentation. Lactate, succinate, acetate, hydrogen, formate, and H_2_S were further metabolised by some community members (Figure 4). The predicted identity of fermentation end-products and other by-products of metabolism was found to be in line with literature information [64, 65, 86]. For example, the formation of lactate was observed for *Lactobacillus acidophilus* and *Bifidobacterium longum*, and butyrate was released by known butyrate producers, i.e. *Faecalibacterium prausnitzii*, *Anaerobutyricum hallii*, *Clostridium perfringens*, and *Coprococcus* spp.. Especially the main products of mixed acid fermentation (acetate, formate, hydrogen, ethanol) were predicted for most members of the community which is in agreement with what is known about common metabolic end products of many gut-dwelling microorganisms [86]. Interestingly, for *Faecalibacterium prausnitzii* no acetate production is reported [86], which was also observed in our simulations. Moreover, H_2_S was correctly predicted to be produced by *Desulfovibrio desulfuricans*. In general, the anaerobic oxidation of fatty acids is not favored by the gut environment because the host competes for the uptake of butyrate, propionate, and acetate, which serve as energy source for colonic epithelial cells and are involved in many host functions [87]. Therefore, the gut community lacks syntrophic organisms which are able to anaerobically degrade butyrate [88]. In agreement with this, we found no microbial uptake of butyrate in the community simulation. In contrast, lactate was predicted to be produced and consumed by distinct community members. We found utilisation of lactate by *Coprococcus comes*, *Megasphaera elsdenii*, and *Veillonella dispar*, which is a known feature of these organisms [64]. In addition, succinate was correctly predicted to be used by *Bacteroides* species [86]. The formation of methane is known to be limited to methanogenic archaea, and thus *Methanosarcina barkeri* produced methane from acetate and hydrogen during our simulations.

**Figure 4:**
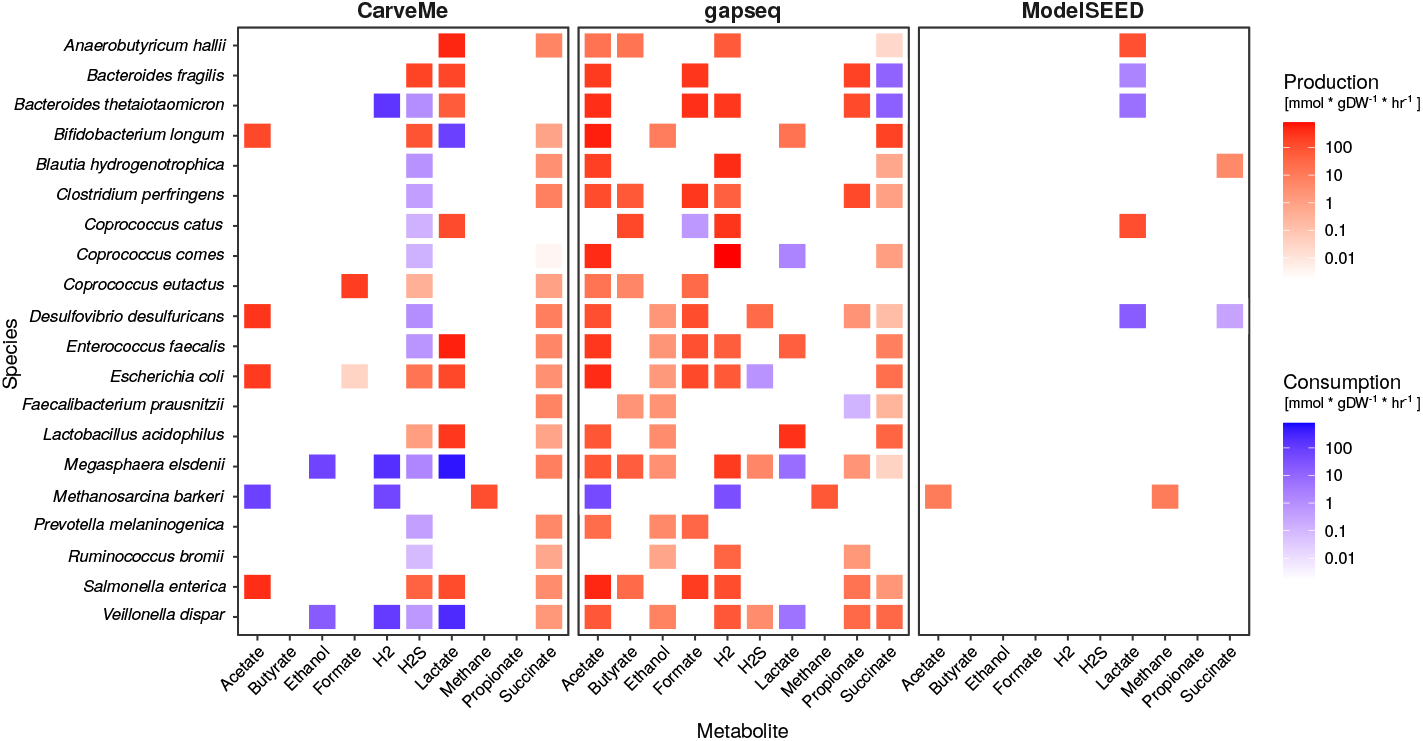
Predicted metabolic products and food web of a microbial community. The metabolism of a community consisting of 19 bacterial species commonly found in the human gut and one archaeon (*Methanosarcina barkeri*) was predicted using BacArena [68]. All bacterial models were reconstructed by CarveMe, `gapseq`, or ModelSEED; with the exception of *M. barkeri* for which a published and manually curated model [66] was used.

For comparison, the community simulation were also performed using models reconstructed with CarveMe and ModelSEED (Figure 4). In both cases, most of the above-mentioned known metabolic cross-feeding interactions and end-products were not predicted, for instance the production of the short chain fatty acids butyrate and propionate was missing. In summary, `gapseq` models were able to recapitulate the major interactions, which are described for microbial communities in the human gut. The overall consumption pattern and individual microbial contributions were found to be in agreement with literature data. Taken together, the community simulation results illustrate the capacity of `gapseq` to construct predictive models for complex metabolic interaction networks comprising several different species.

### 2.7 Pathway prediction of soil and gut microorganisms

To demonstrate the pathway prediction capabilities of `gapseq`, we analysed two communities of soil and gut microorganisms comprising 922 and 822 organisms, repectively. The two communities could be separated from each other by differences in energy metabolism (Principal component analysis, Figure 5A). Here, most variance was explained by subsystems of pathways that are involved in chemoautotrophic, respiratory, and fermentative processes including hydrogen production. Out of 128 energy pathways, the presence of 40 pathways differed significantly (Kolmogorov-Smirnov test, *P* < 0.05) between soil and gut microorganisms and could be categorised into 12 subsystems (Figure 5B). In total, gut microorganisms showed less variety in energy pathways than soil microorganisms. Only pathways relevant for the formation of acetate, hydrogen, and lactate were predicted to be enriched. In the case of all other energy subsystems, more pathways were predicted for soil organisms, most prominently pathways relevant for aerobic and anaerobic respiration as well as the tricarboxylic acid cycle (TCA). In summary, members of the soil community showed a more versatile energy metabolisms, which potentially indicates a higher energetic specialisation of gut microbes. This sample application demonstrates how `gapseq` can facilitate the characterisation and comparison of microbial communities based on the analysis of the presence and absence of specific metabolic pathways.

**Figure 5:**
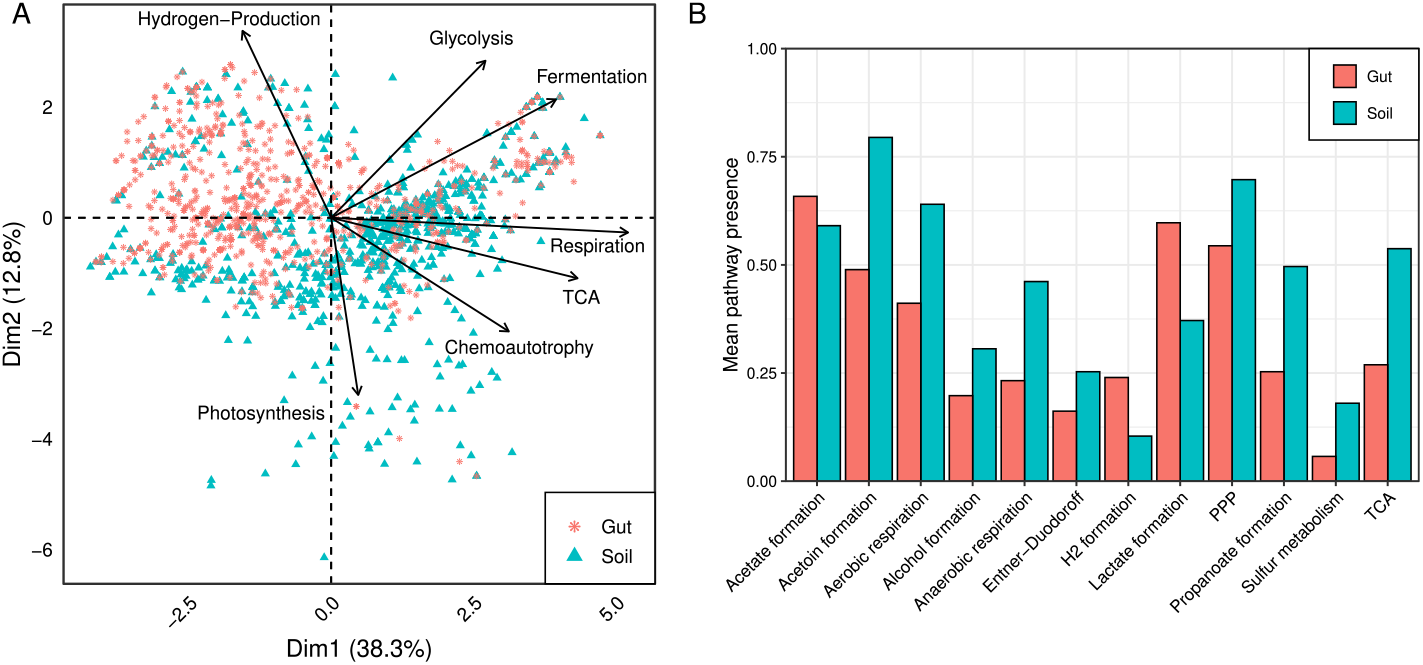
Comparison of energy metabolism between soil and gut community. A) A PCA plot with the first two dimension explaining more than 50% of the variance. Selection of subsystems from energy metabolism with highest quality and impact are shown. B) List of subsystems of energy metabolism that differ significantly in frequency between members of the soil and gut community (TCA: tricarboxylic acid cycle, PPP: Pentose phosphate pathway).

### 2.8 Model reconstructions for metagenomic assemblies

Genome-scale metabolic models can also be reconstructed on the basis of species-level genome bins (SGBs, [69]) assembled from shotgun metagenomic sequencing reads. Yet, genome assemblies from metagenomic material are more prone to errors, fragmentation, and sequence gaps than assemblies of isolated genomes [89], which can potentially cause gaps in the metabolic network reconstructions. We tested whether `gapseq` is able to identify and fill such gaps by comparing the models reconstructed for 127 SGBs from the human microbiome[69] to corresponding models of closely-related reference genomes that were assembled from DNA-sequencing of pure cultures (Figure S2).

As expected, we found a strong positive correlation between the SGBs’ genome completion and their model similarity to their respective reference models (Spearman’s rank correlation, n = 127, *P* < 10^*−*9^). To estimate the quantitative effect of genome completion on the model similarity, a logarithmic function (*y*(*x*) = *c* + *b* ∗ *log*(*x*)) was fitted to the data (*R*^2^ = 0.71, Figure S2). The fitted model indicated, that `gapseq` is able to reconstruct the underlying metabolic network of an organism even on the basis of incomplete and fragmented genomes. For instance, `gapseq` was on average able to recover 90% of the enzymatic reactions that are found in the reference models for SGBs with a predicted genome completion of only 80% (Figure S2).

### 2.9 Summary of validation tests

For each validation approach, predictions were compared to experimental data obtained from databases and literature to calculate prediction performance scores. The overall accuracy (proportion all correct prediction in relation to all predictions made) of model predictions with experimental data was 66% (CarveMe), 70% (ModelSEED), and 81% (`gapseq`)(Table 1). Sensitivity measures the proportion of correctly predicted positives, whereas specificity accounts for the accurate prediction of negatives. All approaches showed a high specificity > 0.7 with highest values for fermentation product and gene essentiality tests. Notably, `gapseq` showed the highest sensitivity over all tests (Figure 6). In summary, `gapseq` outperformed other methods in terms of accuracy and sensitivity while showing similar specificity.

**Figure 6:**
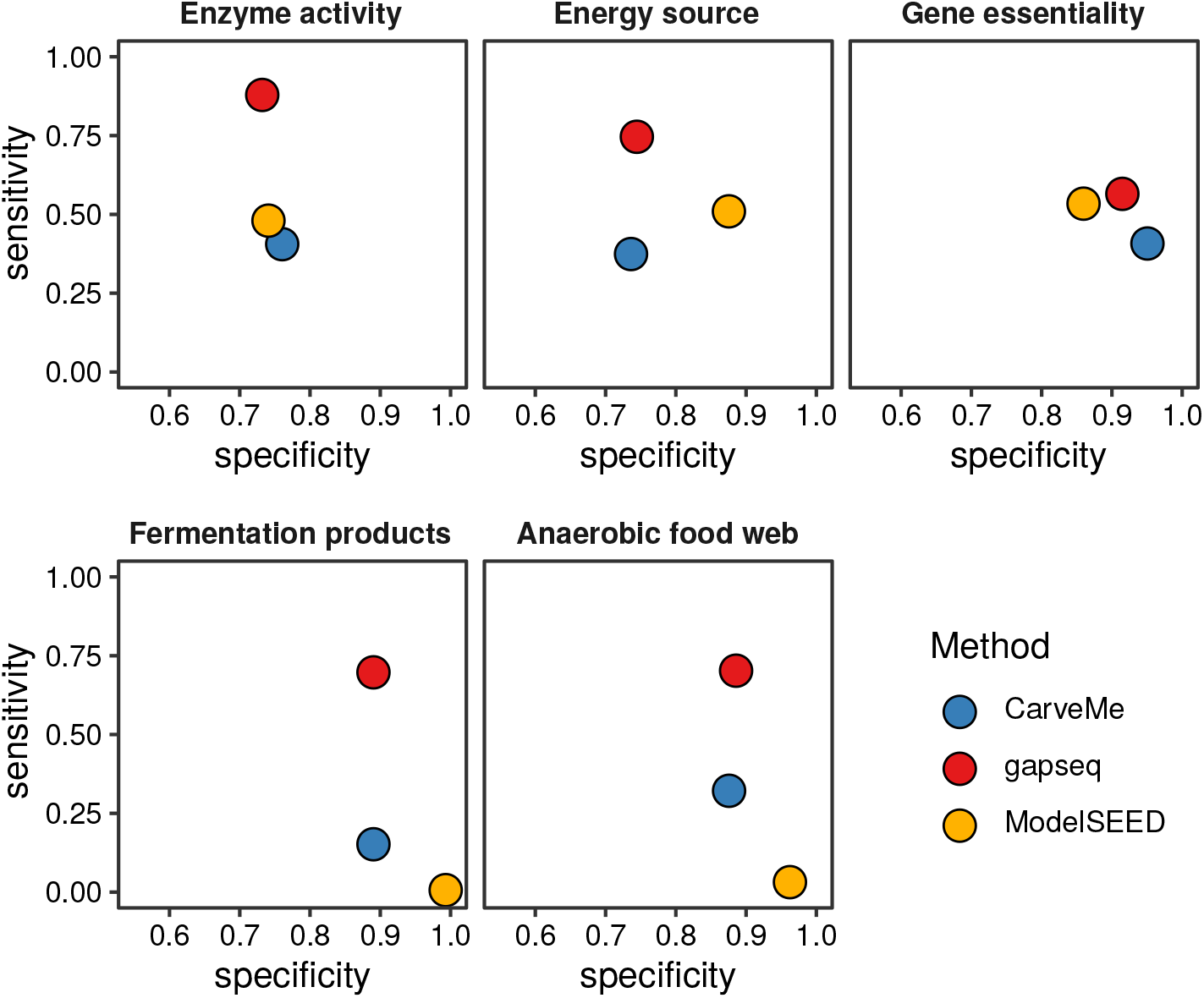
Summary of different validation tests. The specificity and sensitivity for all compared methods are shown. This includes results from benchmarks concerning enzyme activities, energy sources, fermentation products, gene essentiality, and metabolite production/consumption in an anaerobic food web.

**Table 1:** Summary of different methods that were compared in this work. Accuracy, sensitivity, and specificity scores are based on 14, 895 tested phenotypes including energy sources, enzyme activity, fermentation products, gene essentiality, and anaerobic food web structure predictions.

| Metric | CarveMe | gapseq | ModelSEED |
| --- | --- | --- | --- |
| *Implementation* |  |  |  |
| Infrastructure | local | local | web service |
| Input (FASTA file) | protein | nucleotide | nucleotide |
| Programming languages | python | shell script, R | perl/javascript |
| Gap-fill solver | CPLEX | GLPK/CPLEX | not needed* |
| Gap-fill problem formulation | MILP | LP | MILP |
| *Performance* |  |  |  |
| Accuracy | 0.66 | 0.81 | 0.70 |
| Sensitivity | 0.34 | 0.73 | 0.32 |
| Specificity | 0.84 | 0.83 | 0.88 |
| Model file quality** | 0.32 ± 0.006 | 0.78 ± 0.004 | 0.39 ± 0.016 |

\* Solver runs on ModelSEED server. No local solver is required.
\*\* MEMOTE total score ( ± SD).

## 3 Discussion

Here, we introduced `gapseq` - a new tool for metabolic pathway analysis and genome-scale metabolic network reconstruction. The novelty of `gapseq` lies in the combination of (i) a novel reaction prediction that is based both on genomic sequence homology as well as pathway topology, (ii) a profound curation of the reaction and transporter database to prevent thermodynamically infeasible reaction cycles, and (iii) a reaction evidence score-oriented gap-filling algorithm. In order to scrutinise `gapseq` metabolic models, we compared the models’ network structures and predictions with large-scale experimental data sets, which were retrieved from publicly available databases. Furthermore, the ability of `gapseq` to predict bacterial phenotypes was compared to two other commonly used automatic reconstruction methods, namely, CarveMe [39] and ModelSEED [24] (Table 1). ModelSEED is also implemented in the KBASE online software platform [90].

### Crucial large-scale benchmarking of metabolic models

The quality of genome-scale metabolic networks can be assessed by comparing model predictions with experimental physiological data. The protocol by Thiele and Palsson (2010) for the reconstruction of genome-scale metabolic networks recommends the quality assessment and manual network curation using data for (i) known secretion products (e.g. fermentation end-products), (ii) single-gene deletion mutant growth phenotypes (i.e. gene essentiality), and (iii) the utilisation of carbon/energy sources [20]. Tools for the automatic reconstruction of metabolic networks should also make use of such physiological data whenever available for benchmarking. Here, we tested our `gapseq` approach on the basis of all three recommended phenotypic data and compared the performance with CarveMe and ModelSEED. Additionally, we included two novel benchmark tests: The comparison of model predictions with (iv) the activity of specific enzymes known from experimental studies [49] and (v) metabolic interactions (food web) among microorganisms in a multi-species community within an anaerobic environment. Across all five benchmark tests, we could show that `gapseq` outperformed CarveMe and ModelSEED in terms of sensitivity while achieving specificity scores that are comparable to the other two tools (Figure 6).

Publicly available genome sequences of microorganisms, which can be subject for automated metabolic network reconstruction are massively increasing in number due to continuing advances in high-quality and high-throughput sequencing technologies [18]. This development is further fueled by the the increasing number of genome assemblies from metagenomic material [91]. In contrast, standardised phenotypic data for microorganisms remains a bottleneck for the validation of automated metabolic network reconstruction pipelines such as `gapseq`. As consequence, it is crucial for the future development of automated network reconstruction software to include possibly all available phenotypic data for benchmarking, especially data from non-model organisms. To benchmark `gapseq` in in relation to CarveMe and ModelSEED using phenotypic data from mainly non-model organisms, we retrieved phenotypic data of enzyme activity for more than 3, 000 organisms and carbon source utilisation for more than 500 organisms from online databases, which is, to our knowledge, the yet largest phenotypic data set used for validation of automatically reconstructed metabolic networks. In this validation approach `gapseq` achieved the highest prediction accuracy among all three tools tested (Figure 1).

Hence, those results suggest that `gapseq` is a powerful new tool for the automated reconstruction of genome-scale metabolic network models. Moreover, the underlying reference protein sequences as well as the pathway database can readily be updated using online resources, which makes `gapseq` flexible to include future developments and findings in microbial metabolic physiology.

### Automated network reconstructions for community modelling

While single organisms can be considered as the building blocks of microbial communities, individual metabolic models of organisms are the building blocks of *in silico* microbial community simulations. Therefore, genome-scale metabolic models are increasingly applied to predict the function of multi-species microbial communities [61, 92, 93]. To correctly infer metabolic interaction networks between different organisms, it is important that individual models accurately predict nutrient utilisation (e.g. carbon source) and metabolic end-products (e.g. fermentation products). In this study, the benchmarks for carbon source utilisation and fermentation end-product identity indicated that `gapseq` has the highest prediction performance compared to other reconstruction tools (Figure 1 and Figure 3).

To illustrate the applicability of `gapseq`-reconstructed metabolic models for the simulation of multi-species community metabolism, we generated models for microbial strains from the human gut microbiota and simulated their growth in a shared environment. Without further curation, the community simulation reproduced all important hallmarks of intestinal anaerobic food webs [64, 86]. Above all, short chain fatty acids (SCFA) were predicted to be the primary end products of fermentation. This prediction is important to represent intestinal metabolism, because SCFA are crucially involved in host physiology by affecting regulatory response in intestinal and immune cells [94, 95]. Furthermore, the simulation accurately predicted the exchange of metabolites between different members of the microbial community (Figure 4). Cross-feeding of metabolites and the formation of anaerobic food chains have been associated with a healthy microbiome [9, 96]. For instance, the cross-feeding of lactate has been reported to be vital for the early establishment of a healthy gut microbiota in infants [96]. Accordingly we observed the exchange of lactate between different bacterial species in the community simulations (Figure 4) and involved known lactate producers (e.g. *Enterococcus faecalis*) and consumers (e.g. *Megasphaera elsdenii*). This example illustrates that we are able to predict key features of the anaerobic food-web within the gastrointestinal microbiota using `gapseq` models. In addition to the ability to accurately model metabolic processes within existing microbial communities, `gapseq` will further promote the potential of metabolic modelling to predict how complex microbial communities can be modulated by targeted interventions. Specific interventions, which could for instance be predicted, are the introduction of new species to the community (i.e. probiotics) or microbiome-modulating compounds (prebiotics) to the environment. Predictions of potential intervention strategies that target the microbiome are of vast relevance for biomedical research. Furthermore, metabolic interactions between microbiome members are difficult to detect *in vivo* due to the simultaneous production and uptake of metabolites. Thus, *in silico* predictions of metabolite cross-feeding interactions are highly valuable for hypothesis generation about the function and dynamics of microbial communities.

Taken together, the results obtained with `gapseq` suggest, that metabolic models which are reconstructed using `gapseq` are promising starting points to construct ecosystem-scale models of inter-species biochemical processes and to predict targeted strategies to modulate microbiome structure and function.

### Pathway analysis of microbial communities

The construction of genome-scale metabolic models is based on metabolic networks that are inferred from genomic sequences in the context of biochemical databases [20]. Although, the reconstruction of metabolic networks is closely related to the prediction of metabolic pathways, metabolic modelling and pathway analysis are often treated separately [97]. In `gapseq`, the prediction of metabolic pathways is intrinsically tied to the reconstruction of metabolic networks and gap-filling. In addition, reaction, transporter, and pathway predictions can also be used to evaluate the functional capacities of microorganisms without the need of metabolic modelling. As an example for metabolic pathway analysis, we compared the predicted energy metabolism of two large microbial communities that occur in soil and the human gut. We could show that the predicted distribution of pathways differ between both communities based on the habitat, which usually accommodates the members of the respective community. Gut microorganisms showed a less versatile energy metabolism and a specialisation towards fermentation pathways, which lead to the formation of acetate, hydrogen, and lactate. Variations in pathways distributions between both communities may be explained by distinct evolutionary histories. The habitat of the diverse group of soil microorganisms more likely represents an open ecosystem, whereas the gut microbiome is directly constraint by a multi-cellular host that potentially affect microbial phenotypic traits [98]. In general, metabolic modelling should be accompanied by the analysis of pathways based on statistical methods [97] to compensate for additional assumptions, which are introduced in constraint-based metabolic flux modelling [4].

### Limitations and outlook

`gapseq` requires 1-2h for the reconstruction of a single model, whereas ModelSEED and CarveMe operate faster (10min) on a standard desktop computer. Nonetheless, CarveMe needs as input gene sequences (protein or nucleotide), which has to be predicted first, and ModelSEED works as a web service, which can complicate the handling of large-scale reconstruction projects. In `gapseq`, pathways were predicted based on topology and sequence homology searches. However, the assignment of enzymatic function from sequence comparisons has been shown to potentially miss protein domain structures and thus can cause false annotations [99, 100]. In addition, `gapseq` uses many resources to find potential sequences for reactions in pathway databases. Together this might explain why although `gapseq` performed better than other methods on predicting positive phenotypes (function present), it went head to head with regard to negative phenotype predictions (function not present). CarveMe takes a different approach when inferring function by taking care of functional regions (protein domains) [101], resulting in orthologous groups [102], which results in a slightly better specificity (true negative phenotype predictions) in benchmarks (Figure 6). Future developments of `gapseq` will address orthologous groups by using multiple inference methods. Furthermore, the integration of functional predictions coming from phylogenetic inference without the need of genomic sequences [103] might also be promising for further developments of `gapseq`.

## Conclusion

We provide a new software tool called `gapseq` that is suitable for metabolic network analysis and metabolic model reconstruction. To enhance phenotype predictions, `gapseq` employs various data sources and a novel gap-filling procedure that reduces the impact of arbitrary growth medium requirements. We further brought together the so far largest benchmarking of genome-scale metabolic models, in which `gapseq` outperformed comparable alternative tools. With the increased model quality of automated network reconstructions, `gapseq` will provide new insights into the metabolic phenotypes of non-model and yet-uncultured bacteria whose genomes are assembled from metagenomic material. In this way, the models and their simulations allow predictions on the organisms’ ecological role in their natural environments. Taken together, we consider `gapseq` as important contribution to the modelling of microbial communities in the age of the microbiome.

## 4 Methods

### 4.1 Program overview & source code availability

The source code is accessible and maintained at https://github.com/jotech/gapseq. The program is called by `./gapseq`, which is a wrapper script for the main modules. Important program calls are `./gapseq find` (pathway and reaction finder), `./gapseq find-transport` (transporter detection), `./gapseq draft` (draft model creation), `./gapseq fill` (gap-filling), or `./gapseq doall` to perform all in line. When ever necessary, method sections directly refer to config, data and source code files from the `gapseq` package, which contains the main sub-directory `src/` with source code files and `dat/`, which contains databases and also the sequence files in `dat/seq/`. Figure 7 shows an overview of the different `gapseq` modules.

**Figure 7:**
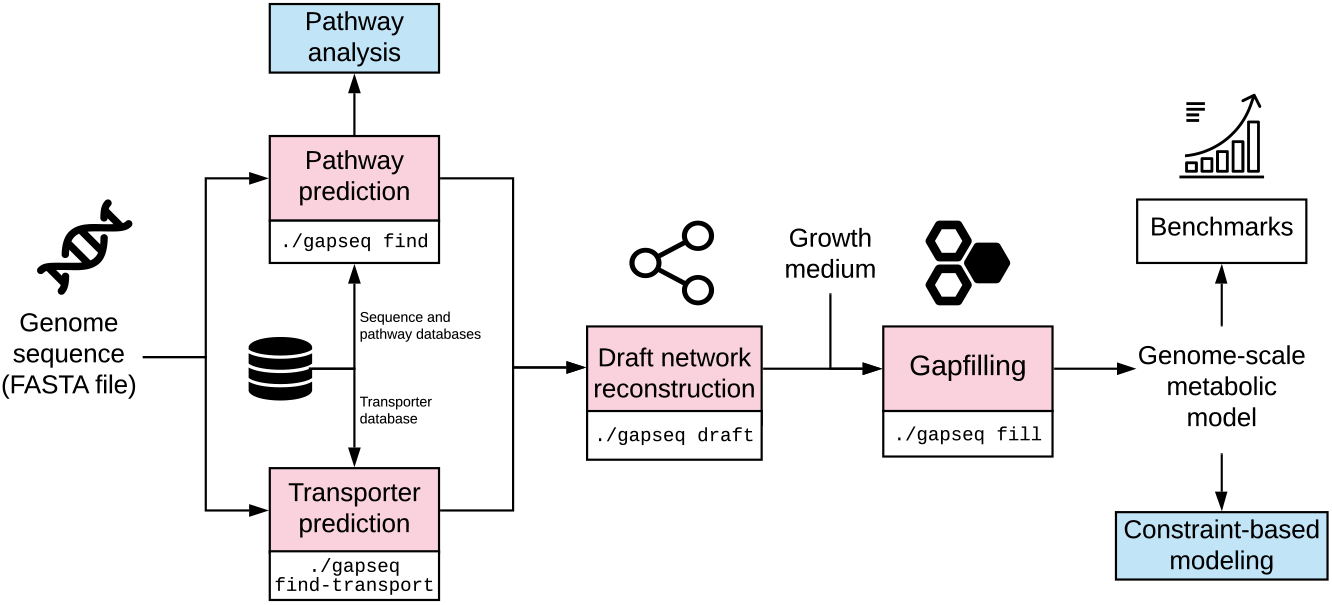
Chart showing the main components and workflow of gapseq. Free icons were used from https://www.flaticon.com (creators: Freepik, Gregor Cresnar, Freepik, Smashicons).

### 4.2 Pathway and sequence databases

Pathways are considered as a list of reactions with enzyme names and EC numbers. Pathway definition were obtained from MetaCyc [27], KEGG [28], and ModelSEED [24]. For MetaCyc, PathwayTools [29] was used in combination with Python-Cyc to obtain pathway definitions [30] (`src/meta2pwy.py`). Information on Kegg pathways were retrieved directly from the KEGG homepage: reactions (http://rest.kegg.jp/list/reaction), and EC numbers (http://rest.kegg.jp/link/pathway/ec) and further processed (`src/kegg_pwy.R`). In case of ModelSEED, subsystem definition were obtained from the homepage: http://modelseed.org/genomes/Annotations (`src/seed_pwy.R`). In addition, manual defined and revised pathways are stored in the file `dat/custom_pwy.tbl`.

Sequence data needed for pathway prediction were downloaded from UniProt [31] for each reaction identified by EC number, enzyme name, or gene name. Both reviewed and unreviewed sequences are considered and stored as clustered UniPac sequences (`src/uniprot.sh`). To increase the sequence pool for a given reaction, alternative EC numbers from BRENDA [32] and from the Enzyme Nomenclature Committee https://www.qmul.ac.uk/sbcs/iubmb/enzyme/ are integrated (`src/altec.R`, `dat/brenda_ec.csv`).

### 4.3 Pathway prediction

For each pathway selected from a pathway database (MetaCyc, KEGG, ModelSEED, custom), `gapseq` searches for sequence evidence and a pathway is defined as present if enough of its reactions were found to have sequence evidence. In more detail, sequence data (section 4.2) is used for homology search by *tblastn* [33] with the protein sequence as query and the genome as database. By default, a bitscore *≥* 200 and a coverage of at least 75% is needed for a match. For certain reactions, the user can define additional criteria, for example an identity of *≥* 75% (`dat/exception.tbl`). In case of protein complexes with subunits, a more complex procedure is followed (section 4.4). Spontaneous reactions, which do not need an enzyme, were set to be present in any case. In general, a pathway is considered to be present if at least 80% of the reactions are found (completenessCutoffNoHints threshold). This pathway completeness threshold is lowered for pathways in following cases:

1. If the pathway contains key reactions, as it is defined for some pathways in MetaCyc, and all key reactions are found, then `completenessCutoff` of the total reactions needed to be found. We used a value of 2/3 for this threshold.
2. In cases in which no sequence data is available for specific reactions, the status of the reactions is set to “vague” and these reactions do not count as missing if they account for less than `vagueCutoff` of the total reactions of a pathway. We used a value of 1/3 for this threshold.

The pathway prediction algorithm is implemented in the bash shell script `src/gapseq_find.sh`, which uses GNU parallel [34] and fastaindex/fastafetch from exonerate [35].

### 4.4 Protein complex prediction

A problem with automatic sequence download for reactions (as FASTA files) comes with protein complexes, for which a single blast hit may be not sufficient to predict enzyme presence. In `gapseq`, subunits are detected by text matching in the FASTA headers. Search terms are: “subunit”, “chain”, “polypeptide”, “component”, and different numbering systems (roman, arabic, greek) are homogenised. To avoid artifacts in text matching, subunits that occur less than five times in the sequence file are not considered, and in cases in which a subunit occurs almost exclusively (*≥* 66%) the other entries are not taken into account. All FASTA entries, which could not matched by text mining, or which are excluded because of the coverage, are labeled ‘undefined subunit’ and do not add to the total amount of subunits. For each recognised subunit, a blast search is done. A protein complex counts as present if more than 50% of the subunits could be found, whereby the presence of ‘undefined subunits’ tip the balance if exactly 50% of the subunits were found. The text matching with regular expressions is done with R’s stringr [36] and biostrings [37] as defined in `src/complex_detection.R`. The script is called from within the shell script `src/gapseq_find.sh`.

### 4.5 Transporter prediction

For transporter search, sequence data from the Transporter Classification Database is employed [38]. In addition, manual defined sequences can be defined in `dat/seq/transporter.fasta`. The sequence set is reduced to a subset of transporters that involve metabolites known to be produced or consumed by microorganisms (`dat/sub2pwy.csv`). Subsequently, the genome is queried by the reduced sequences using *tblastn* [33]. For each hit (default cutoffs: bitscore *≥* 200 and coverage *≥* 75%), the transporter type (1. Channels and pores, 2. Electrochemical potential-driven transporter, 3. Primary active transporters, 4. Group translocators) is determined using the TC number mentioned in the FASTA header. A suitable candidate reaction is searched in the reaction database. If there is a hit for a transporter of a substance but no candidate reaction for the respective transporter type can be found, then other transporter types are considered. The transporter search is done by the shell script `src/transporter.sh` that uses GNU parallel [34] and fastaindex/fastafetch from exonerate [35].

Candidate transporters are selected from the reaction database by transporter type and substance name. This is done by text search and is currently implemented only for the ModelSEED namespace. From the ModelSEED reaction database all reaction with the flag *is transport* = 1 are taken and the transporter type is predicted by keywords: “channel”, “pore” (1. Channels and pores); “uniport”, “symport”, “antiport”, “permease”, “gradient” (2. Electrochemical potential-driven transporters); “ABC”, “ATPase”, “ATP”(3. Primary active transporters); “PTS” (4. Group translocators). If no transporter type could be identified by keywords, additional string matching is done for ATPases, proton/sodium antiporter, and PTS by considering the stoichiometry of the involved metabolites. The transported substance is identified as the substance that occurs on both sides of the reaction. In addition, reactions from the reaction database can be linked manually to substances and transporter types (`dat/seed_transporter custom.tbl`). The text matching with regular expressions is done with stringr [36] (`src/seed_transporter.R`).

### 4.6 Biochemistry database curation and construction of universal metabolic model

For the construction of genome-scale metabolic network models, `gapseq` uses a reactions and metabolite database that is derived from the ModelSEED database [24] as from January 2018. In addition, 30 new reactions and 2 new metabolites were introduced to the `gapseq` biochemistry database (see suppl. table S1). All reactions and metabolites from the database were included for the construction of a full universal metabolic network model; an approach that is also used in CarveMe [39]. We curated the underlying biochemistry database in order to correct inconsistencies in reaction stoichiometries and reversibilities. Inconsistencies were identified by optimising the universal network model for ATP-production without any nutritional input to the model using flux balance analysis. In case of ATP-production, the flux distributions of such thermodynamically infeasible reaction cycles were investigated by cross-checking the involved reactions with literature information, the BRENDA database for enzymes [32], and the MetaCyc database [27]. Stochiometries and reversibilities of erroneous reactions were corrected accordingly. This curation procedure was repeated until no theromodynamically infeasible and ATP-generating reaction cycles were observed.

Hits from the pathway prediction (4.3) and transporter prediction (4.5) are mapped to the `gapseq` reaction database using different common identifiers. A majority of reactions are directly matched via their corresponding Enzyme Commission (EC) system identifier [40] and Transporter Classification (TC) system identifier [38], respectively. For this mapping, also alternative EC-numbers for enzymatic reactions as defined in the BRENDA database [32] are considered. Moreover, the databases used for pathway and transporter predictions often provide cross-links to the reaction’s KEGG ID, which is also assigned to most reactions in the `gapseq` database and used to match reactions. Additionally, the MNXref database [41] provides cross links between several biochemistry databases, which `gapseq` also utilises to translate hits from the pathway predictions to model reactions. Finally, a manual translation of enzyme names to model reactions is done for some reactions, which we identified as important reactions but which failed to match between the pathway databases (4.3) and the `gapseq` model reactions using other reaction identifiers (`dat/seed_Enzyme_Name_Reactions_Aliases.tsv`). The overall mapping is done by the function `getDBhit()` as defined in `./src/gapseq_find.sh`.

### 4.7 Model draft generation

A draft genome-scale metabolic model is constructed based on the results from the pathway and transporter predictions (see above). A reaction is added to the draft model if the corresponding enzyme/transporter was directly found or if the pathway was predicted to be present (i.e. due to pathway completeness and key enzymes) in which the reaction participates. Additionally, spontaneous reactions as defined in the MetaCyc database as well as transport reaction of compounds, which are know to be able to cross cell membranes by means of diffusion (e.g. *H*_2_), are directly added to every draft model. As part of the draft model construction `gapseq` adds a biomass reaction to the network that aims to describe the composition of molecular constituents that the organism needs to produce in order to form 1 g dry weight (1 gDW) of bacterial biomass. `gapseq` uses the biomass composition definition from the ModelSEED database for Gram-positive (`dat/seed_biomass.DT_gramPos.tsv`) and Gram-negative bacteria (`dat/seed_biomass.DT_gramNeg.tsv`). If no Gram-staining property is specified by the user, `gapseq` predicts the Gram-staining-dependent biomass reactions by finding the closest 16S-rRNA-gene neighbor using a blastn search against reference 16S-rRNA gene sequences from 4647 bacterial species with known Gram-staining properties that are obtained from the PROTRAITS database [42]. The model draft generation is done by the R script `src/generate_GSdraft.R`.

### 4.8 Gap-filling algorithm

`gapseq` provides a gap-filling algorithm that adds reactions to the model in order to enable biomass production (i.e. growth) and likely anabolic and catabolic capabilities. The algorithm uses the alignment statistics (i.e. the bitscore) from the pathway- and transporter prediction steps of `gapseq` (see above) to preferentially add reactions to the network, which have the highest genetic evidence. This approach is especially relevant in cases where the sequence similarity to known enzyme-coding reference genes was close to but did not reach the cutoff value *b*, which is required for a reaction to be included directly into the draft network. In contrast to the gap-filling algorithms described in previous works [43] and [39], which also use genetic evidence-weighted gap-filling, the gap-filling problem in `gapseq` is not formulated as Mixed Integer Linear Program (MILP) but as Linear Program (LP), and is derived from the parsimonious enzyme usage Flux Balance Analysis (pFBA) algorithm developed by Lewis *et al.*, 2010 [3]. Therefore, the alignment statistics (i.e. bitscore) are translated into weights for the corresponding model reactions and incorporated into the problem formulation:

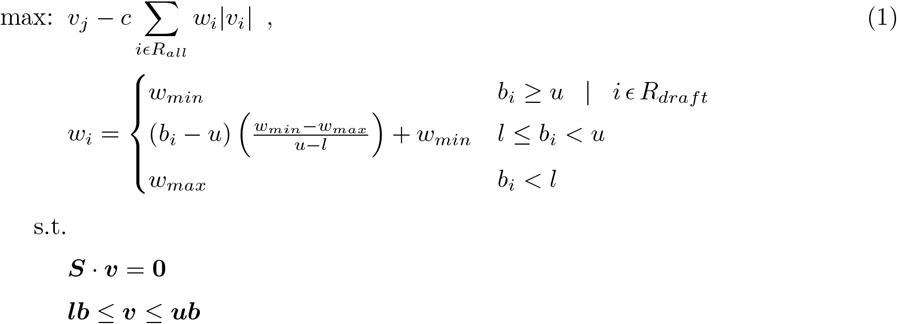

Where *R_all_* is the set of all reaction in the universal model, *R_draft_* are the reactions, which are already part of the draft network before gap-filling, *v_j_* is the flux through the objective reactions (e.g. biomass production), *v_i_* the flux through reaction *i*, *w_i_* the weight for reaction *i*, *v* the flux vector for all reactions, and *c* a scalar factor that determines the contribution of the absolute reduction of weighted fluxes to the overall FBA solution (default: *c* = 0.001). Moreover, a maximum weight value *w_max_* (default: 100) is assigned if the reaction’s highest bitscore is smaller than a threshold *l* (default: 50). A minimum reaction weight *w_min_* (default: 0.005) is assigned to reactions with a bitscore higher than *u* (default: 200) or if the reactions are already part of the draft model. *S* is the stoichiometric matrix and *lb* and *ub* the lower and upper flux bound vectors.

Two other LP-based gap-filling algorithms that incorporate reaction evidence scores have been formulated by Dreyfuss *et al.* (2013) [44] and Medlock *et al.* (2020) [45], respectively. These approaches require a definition of a minimum flux through the biomass reaction to ensure growth. The pFBA-derived LP formulation of `gapseq` (equation 1) includes the flux through the biomass/objective reaction *v_j_* together with the reaction evidence scores in a single objective function. In `gapseq` and following the solution of the LP (1), reactions carrying a flux and which are not part of the draft model are added to the network model. The algorithm is implemented in `src/gapfill4.R`.

### 4.9 Gap-filling of biomass, carbon sources, and fermentation products

Gap-filling of a draft model in `gapseq` requires only for the first step a user-defined growth medium that is ideally known to support growth of the organism of interest *in vivo*. If no growth medium is specified by the user, a complete medium (ALLmed) is chosen by `gapseq` (as done for the large-scale benchmarks of enzyme activity and carbon sources, cf. 4.11, 4.12). A set of common microbial growth media (e.g. LB, TSB, M9) is provided in the `gapseq` software directory `dat/medium/`. In addition, the user can provide a custom growth medium definition. The above described gap-filling algorithm is used to improve the generated draft model in four steps.

1. **Biomass production**: To ensure that the model is able to produce biomass under the given nutritional input (medium) the gap-filling algorithm is applied while the objective is defined as the flux through the biomass reaction. This step will add all missing reactions that are essential for *in silico* growth.
2. **Individual biomass components**: It is checked whether the model supports the biosynthesis of biomass components. Therefore, model is re-constrained to a M9-like minimal medium with a carbon source for which an exchange reactions is found (default: glucose if available). The objective function is set to the production of one biomass component at a time and the gap-fill algorithm is performed. This gap-filling step is repeated for each biomass component metabolite twice, with and without oxygen to potentially allow aerobic and anaerobic growth for facultative anaerobe species.
3. **Alternative energy sources**: `gapseq` attempts to gap-fill likely metabolic pathways, which enable the utilisation of alternative energy sources, which might not be part of the defined growth medium from step (1). To this end, the model is re-constrained to a M9-like minimal medium containing a single carbon source of interest at the time. As objective function, the summed flux of artificial reactions that accept electrons from the electron carriers ubiquinol, menaquinol, or NADH is defined. This test can be considered as an *in silico* simulation of the commonly used BIOLOG carbon source utilisation test arrays [46] in which the colometric effect is coupled to a dehydrogenase [47]. This gap-filling step is performed for all metabolites defined in `dat/sub2pwy.csv`.
4. **Metabolic products**: Finally, the same list of compounds as for step (3), is used to check whether the network can be gap-filled to allow the formation of these metabolites given the original medium. For each compound the gap-filling algorithm is applied with the production of the focal compound as objective function.

While step (1) considers all reaction from the universal model as potential candidate reactions for gap-filling, steps (2-4) allow only the addition of candidate reactions to the model with a corresponding bitscore from the pathway prediction (4.3) higher than a threshold value *b* (default: 50). Thus, these so-termed ‘*core reactions*’ represent only reactions, for which `gapseq` has found genomic sequence or pathway evidence. This approach for steps (2-4) is chosen to avoid the addition of biosynthetic capabilities to the model, which the organism presumably does not possess.

### 4.10 Formal and functional model file testing

The validity of genome-scale metabolic model files was checked with MEMOTE (0.10.2) [48]. For all models used in the anaerobic food web (4.16), the total MEMOTE score was computed for the respective SBML-Model files. MEMOTE was executed using the parameter `--skip test_find_metabolites_not_produced_with_open_bounds` and `--skip test_find_metabolites_not_consumed_with_open_bounds` since these tests do not contribute to the total MEMOTE score but require long computation time.

### 4.11 Validation with enzymatic data (BacDive)

The Bacterial Diversity Metadatabase (BacDive) [49] was used to obtain enzymatic activity data. For this purpose, a list of type strains IDs where downloaded using the advanced search. Afterwards the IDs were used to query the database via the R package BacDiveR (0.9.1) to obtain the data [50]. If the stored data contained non-zero entries for enzymatic activity and if a genome assembly was available on NCBI, the type strain was considered for the validation analysis. The respective genome assemblies were downloaded with `ncbi-genome-download` (https://github.com/kblin/ncbi-genome-download). If multiple genomes were available for one type strain, ‘*representative*’ and ‘*complete*’ (NCBI tags) genomes were preferred and, in case there were still multiple candidate genomes available, the most complete genome was selected. Genome completeness was estimated by employing the software BUSCO (3.0.2) [51]. In total, 3017 type strain genomes were taken as input for ModelSEED (2.5.1), CarveMe (1.2.2), and `gapseq` to create metabolic models. The gap-filling parameters were set to default values for each program, i.e. a complete medium was assumed. The final test whether a reaction activity is covered by a model was done by checking if the corresponding reaction is present in the model. This was done by matching enzymes and reactions via EC numbers. For CarveMe the vmh (https://www.vmh.life) and for ModelSEED and `gapseq` the ModelSEED (http://modelseed.org) reaction database was used to match reactions and EC numbers. For the EC numbers 3.1.3.1, 3.1.3.2, the corresponding reactions were the same, and thus unspecific, so that both EC numbers were not considered for the validation analysis. In general, the enzyme activities in the BacDive database have the form active (”+”) or not active (“−”) but some entries were ambiguous (e.g.: “+/−”). The ambiguous entries were omitted from the analysis.

### 4.12 Validation with carbon sources data (ProTraits)

Data for the validation of carbon source utilisation was obtained from the “atlas of prokaryotic traits” database (ProTraits) [42]. A tab-separated table with binarised predictions with a stringent threshold of precision of *≥* 0.95 were downloaded from http://protraits.irb.hr/data.html. For organisms which had at least one carbon source prediction, the corresponding genome was obtained from NCBI Ref-Seq [52] if available. In cases where a genome assembly was found, it was taken as input for ModelSEED, CarveMe, and `gapseq` to create metabolic models. The number of potential carbon sources was reduced to a subset for which a mapping from substance name to ModelSEED and CarveMe model namespace existed (`dat/sub2pwy.csv`). The tests for D-lyxose were removed because it was listed as all negative in ProTraits and also all compared pipelines predicted no utilisation. The main test whether a carbon source can be used by a model was done in a BIOLOG-like manner as described above (see 4.9). To this end, temporary reactions to recycle reduced electron carriers as carbon source utilisation indicators were added to the respective model. The objective for optimisation was set to maximise the flux through these recycling reactions. The exchange reactions were limited to a minimal medium with minerals and the focal potential carbon source. This theoretical approach tested, whether the model is able to pass electrons from the potential carbon source to electron carrier metabolites. A carbon source was predicted to be able to serve as energy source if the recycle reactions carried a positive flux.

### 4.13 Prediction of gene essentiality

To predict the essentiality of genes we performed *in silico* single gene deletion phenotype analysis for the network reconstructions of *Escherichia coli* str. K-12 substr. MG1655 (RefSeq assembly accession: GCF 000005845.2), *Bacillus sub-tilis* substr. *subtilis* str. 168 (GCF 000789275.1), *Shewanella oneidensis* MR-1 (GCF 000146165.2), *Pseudomonas aeruginosa* PAO1 (GCF 000006765.1), and *Mycoplasma genitalium* G37 (GCF 000027325.1). The analysis was performed on the basis of the models’ Gene-Protein-Reaction (GPR) mappings and according to the protocol by Thiele and Palsson, 2010 [20]. To this end, the contingency tables of predicted growth/no growth phenotypes from the network models and experimentally determined growth phenotypes of gene deletion mutants were constructed. Genes were predicted to be conditionally essential under the given growth environment if the predicted growth rates of the models were below 0.01 hr^−1^. The growth media compositions for growth predictions were defined as M9 with glucose as carbon- and energy source for *E. coli*, lysogeny broth (LB) for *B. subtilis* and *S. oneidensis*, M9 with succinate as carbon and energy-source for *P. aeruginosa*, and a complete medium (all external metabolites available for uptake) for *M. genitalium*. Experimental data for gene essentiality was obtained from [53, 54, 55, 56, 57].

### 4.14 Fermentation product tests

The release of by-products from anaerobic metabolism was predicted using Flux Balance Analysis (FBA) coupled with a minimisation of total flux [58] to avoid fluxes that do not contribute to the objective function of the biomass production. In addition, Flux-Variability-Analysis (FVA) [59] was applied to predict the maximum fermentation product release of individual metabolites across all possible FBA solutions. Metabolites with a positive exchange flux (i.e. outflow) were considered as fermentation products. The analysis was performed for 18 different bacterial organisms, which (1) have a genome assembly available in the RefSeq database [52], (2) are known to grow in anaerobic environments, and (3) for which the fermentation products have been described in the literature based on anaerobic cultivation experiments (suppl. table S2). The gap-filling of the network models using `gapseq`, CarveMe, and ModelSEED as well as the simulations of anaerobic growth were all performed assuming the same growth medium that comprised several organic compounds (i.e. carbohydrates, polyols, nucleotides, amino acids, organic acids) as potential energy sources and nutrients for growth (see media file `dat/media/FT.csv` at the gapseq github repository).

Since the amount of fermentation product release depends on the organism’s growth rate, we normalised the outflow of the individual fermentation products, which has the unit *mmol* ∗ *gDW^−^*^1^ ∗ *hr^−^*^1^, by the predicted growth rate of the respective organism which has the unit *hr^−^*^1^. Thus, we report the amount of fermentation product production in the quantity of the metabolite that is produced per unit of biomass: *mmol* ∗ *gDW^−^*^1^.

### 4.15 Pathway prediction of soil and gut microorganisms

The pathway analysis was done by comparing predicted pathways of soil and gut microorganisms. For this means, genomes were downloaded from a resource of reference soil organisms [60] and gut microbes [61]. The default parameter of `gapseq` were used for pathway prediction. The principal component analysis was done in R using the factoextra package [62]. For predicted pathways for soil and gut microorganisms, it was checked if samples belong to different distributions using a bootstrap version of the Kolmogorov-Smirnov test [63].

### 4.16 Anaerobic food web of the human gut microbiome

Representative bacterial organisms known to be relevant in the human intestinal cross-feeding of metabolites were selected based on the proposed food webs by Louis *et al.*, 2014 [64] and Rivera-Chavez *et al.*, 2015 [65]. The genomes of organisms were obtained from NCBI RefSeq [52] and metabolic models reconstructed using `gapseq`, carveme, and modelseed. A medium containing minerals, vitamins, amino acids, fermentation- and metabolic by-products (namely acetate, formate, lactate, butyrate, propionate, H_2_, CH_4_, ethanol, H_2_S, succinate), and carbohydrates (glucose, fructose, arabinose, ribose, fucose, rhamnose, lactose) was used for gap-filling.

Furthermore, a published model of *Methanosarcina barkeri* was added to the community [66] to represent archaea that are also known to be part of anaerobic food webs [67]. All organisms of the modeled community and their respective genome assembly accession numbers are listed in supplementary table S3. All metabolic models were then simulated with BacArena [68] by using the described medium but without the fermentation and by-products, plus sulfite and 4-aminobenzoate which were needed for growth by the *M. barkeri* model. The community was simulated for five time steps (corresponding to 5 hours simulated time). The analysis of metabolite uptake and production were done after the third time step, for which all organisms were still growing exponentially.

### 4.17 Model reconstructions from metagenomic assemblies

4, 930 species-level genome bins (SGBs) assembled from shotgun metagenome sequencing reads were obtained from the study of Pasolli *et al.*, 2019 [69]. Only those SGBs were considered for further analysis, which were already classified as bacteria on a species-level in the original publication by Pasolli *et al.*. For each SGB, closely related reference assemblies from the RefSeq database [52] were identified by constructing a multi-locus phylogenetic tree using autoMLST (version as of April 7th 2020, [70]). RefSeq assemblies were considered as genomes from the same species-level taxonomic group as the focal SGB if their predicted MASH distance (*D*)[71] were below or equal to 0.05. This threshold was shown before to cluster bacterial genomes at the taxonomic level of species [71]. Only SGBs with 10 or more assigned reference assemblies were considered for further analysis, which yielded in total 127 SGBs. Metabolic models were reconstructed using `gapseq` for each SGB and their 10 closest reference assemblies (Suppl. Table S5).

Next, similarity of SGB models with their respective reference models was calculated using the following metabolic network similarity score *T_SGB_*:

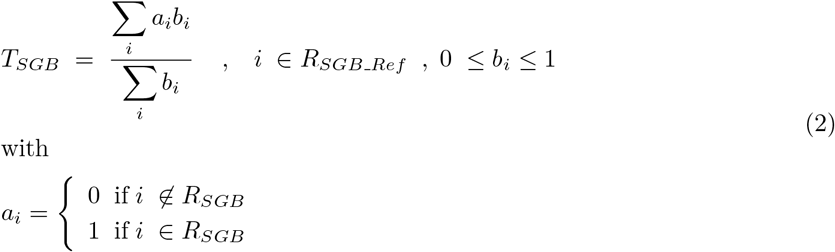

*R_SGB_Ref_* is the union set of reactions with associated genes that are part of the network models reconstructed for the ten reference genome assemblies of the focal SGB. *R_SGB_* is the set of reactions part of the SGB’s model reconstruction. *b_i_* is the frequency of reaction *i* among the ten SGB’s reference models.

Completion of the genome sequence of SGBs was estimated by using BUSCO (version 4.0.6, [51]) using the specific completion score.

### 4.18 Technical details

The pathway prediction part of `gapseq` is implemented as Bash shell script and the metabolic model generation part is written in R. Linear optimisation can be performed with a different solvers (GLPK or CPLEX). Other requirements are exonerate, bedtools, and barrnap. In addition, the following R packages are needed: data.table [72], stringr [73], sybil [74], getopt [75], reshape2 [76], doParallel [77], foreach [78], R.utils [79], stringi [80], glpkAPI [81], and BioStrings [82]. Models can be exported as SBML [83] file using sybilSBML [74] or R data format (RDS) for further analysis in R, for example with sybil [74] or BacArena [68].

## Author’s contributions

JZ, CK, and SW conceptualized `gapseq`. JZ and SW developed the software and did the analysis. JZ, CK, and SW wrote the manuscript.

## Acknowledgements

We thank Martin Sperfeld for fruitful comments and discussions during the developmental phase. The software was thankfully tested by Georgios Marinos, Shan Zhang, and Lena Best.

## Availability of data and materials

`gapseq` is implemented in R and python and is freely available under the GNU General Public License (v3.0) on GitHub (https://github.com/jotech/gapseq/). All results presented in this manuscript were produced using the specific `gapseq` version 1.0 as archived on GitHub. The datasets used for model construction and validation purposes were obtained from publicly available databases and publications as cited at the respective parts of the manuscript.

## Additional Files

Table S1 — New reactions and metabolites added to biochemistry database.

see file: `Table S1.xlsx`

Table S2 — Organisms included in fermentation product validation test.

see file: `Table S2.xlsx`

Table S4 — References for substance production and consumption in anaerobic gut communities (see Supplementary Figure S1).

see file: `Table S4.ods`

Table S5 — 127 Species-level genome bins (SGBs) from Pasolli *et al.*, 2019 [69] and 1270 mapped reference genome assembles from RefSeq.

see file: `Table S5.ods`

**Table S3.** Organisms used in modelling of the anaerobic food web of the human gut microbiome.

| RefSeq Assembly | Organism name | Reconstruction method |
| --- | --- | --- |
| GCF_000173975.1 | *Anaerobutyricum hallii* DSM 3353 | gapseq / modelseed / carveme |
| GCF_000025985.1 | *Bacteroides fragilis* NCTC 9343 | gapseq / modelseed / carveme |
| GCF_001314975.1 | *Bacteroides thetaiotaomicron* | gapseq / modelseed / carveme |
| GCF_000196555.1 | *Bifidobacterium longum* subsp. *longum* JCM 1217 | gapseq / modelseed / carveme |
| GCF_000157975.1 | *Blautia hydrogenotrophica* DSM 10507 | gapseq / modelseed / carveme |
| GCF_000013285.1 | *Clostridium perfringens* ATCC 13124 | gapseq / modelseed / carveme |
| GCF_003434235.1 | *Coprococcus catus* | gapseq / modelseed / carveme |
| GCF_000155875.1 | *Coprococcus comes* ATCC 27758 | gapseq / modelseed / carveme |
| GCF_000154425.1 | *Coprococcus eutactus* ATCC 27759 | gapseq / modelseed / carveme |
| GCF_000189295.2 | *Desulfovibrio desulfuricans* ND132 | gapseq / modelseed / carveme |
| GCF_000391485.2 | *Enterococcus faecalis* EnGen0107 | gapseq / modelseed / carveme |
| GCF_000005845.2 | *Escherichia coli* str. K-12 substr. MG1655 | gapseq / modelseed / carveme |
| GCF_000162015.1 | *Faecalibacterium prausnitzii* A2-165 | gapseq / modelseed / carveme |
| GCF_003047065.1 | *Lactobacillus acidophilus* | gapseq / modelseed / carveme |
| GCF_001304715.1 | *Megasphaera elsdenii* 14-14 | gapseq / modelseed / carveme |
| GCF_000195895.1 | *Methanosarcina barkeri* str. *Fusaro* | manually curated (BiGG-ID: iAF692)<sup>[66]</sup> |
| GCF_000144405.1 | *Prevotella melaninogenica* ATCC 25845 | gapseq / modelseed / carveme |
| GCF_900101355.1 | *Ruminococcus bromii* | gapseq / modelseed / carveme |
| GCF_000006945.2 | *Salmonella enterica* subsp. *enterica* serovar *Typhimurium* str. LT2 | gapseq / modelseed / carveme |
| GCF_900637515.1 | *Veillonella dispar* | gapseq / modelseed / carveme |

**Figure S1.**
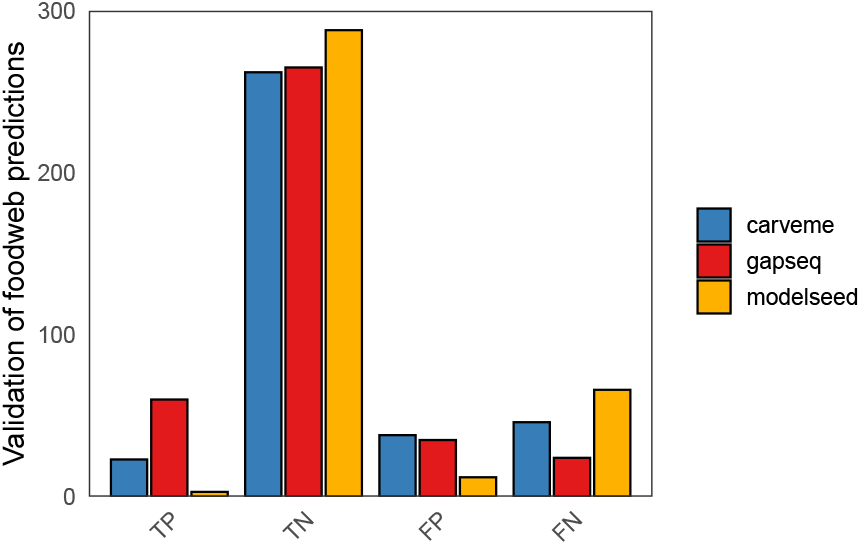
Validation of substance production and consumption in anaerobic gut communities (see Supplementary Table S4).

**Figure S2.**
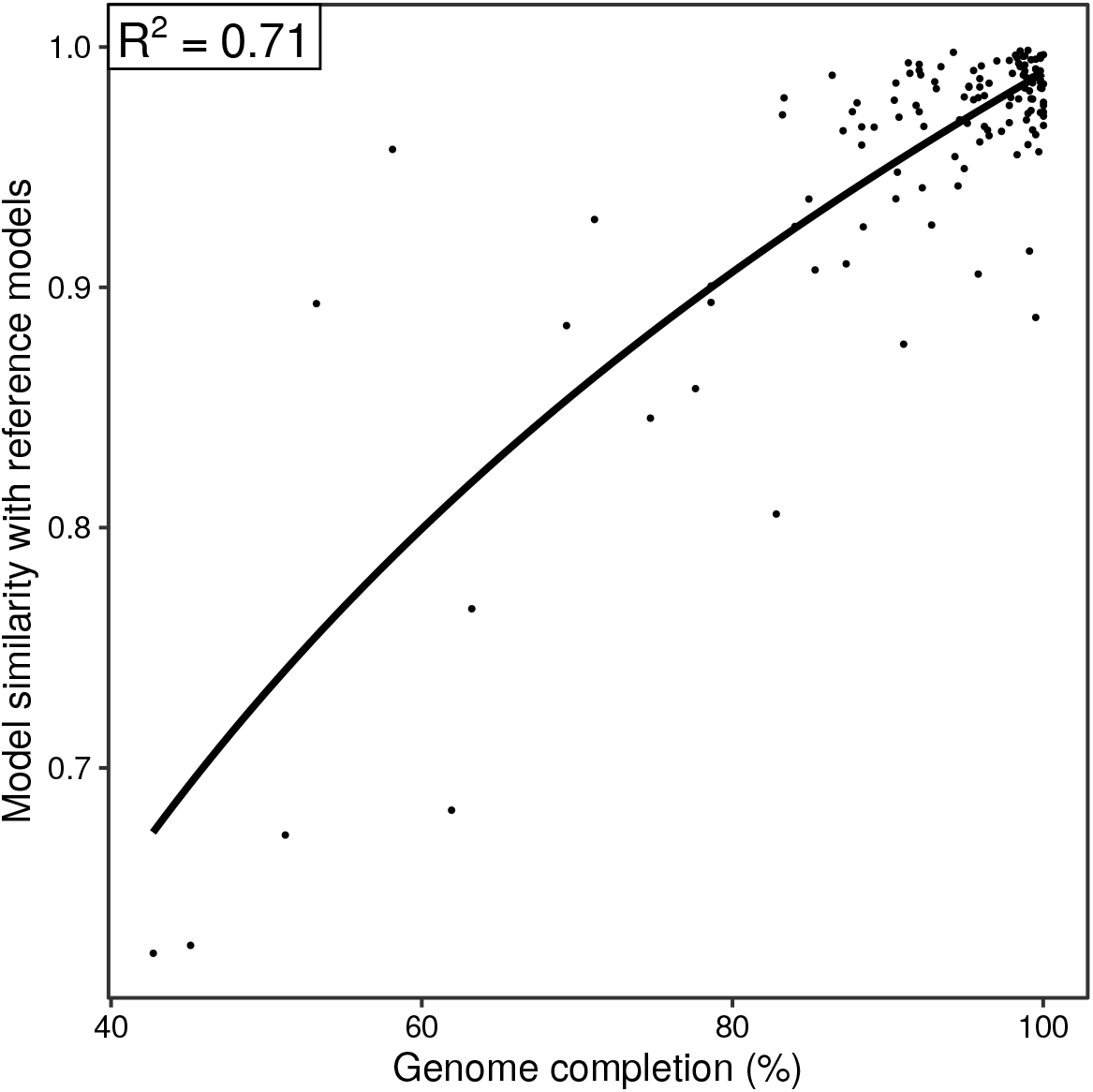
Similarity of gapseq models reconstructed for 127 species-level genome bins (SGBs) from metagenomes compared to models reconstructed for reference genomes (RefSeq Prokaryotic Genomes). The x-axis represents the genome assembly completion of SGBs estimated using the BUSCO software version 4.0.6 [51]. The line shows the result of non-linear regression using a logarithmic function of form *y*(*x*) = *c* + *b* ∗ *log*(*x*). Sequences of SGBs were obtained from Pasolli *et al.*, 2019 [69].

